# Formation, persistence, and breakdown of carrier-set topology in linkage disequilibrium: empirical structure in 1000 Genomes and a two-locus Wright–Fisher model

**DOI:** 10.64898/2026.07.01.735767

**Authors:** Yuki Ichikawa

## Abstract

Linkage disequilibrium between two biallelic loci is usually summarized by scalar association measures such as *r*^2^ and *D*^′^. These measures quantify how visible an allelic association is to a symmetric LD scan, but they do not directly represent the topology of carrier sets: whether the carriers of one variant are contained within, partially overlap with, or are disjoint from the carriers of the other. This distinction is structural. On the haplotype-frequency simplex, carrier-set inclusion corresponds to a boundary face where one haplotype class is absent. In the rare–common regime, a nested rare variant is further constrained by the ceiling *r*^2^ ≤ *p*_*A*_/*p*_*B*_, so that complete carrier-set inclusion can remain nearly invisible to *r*^2^.

Here, as a companion to the Fisher-geometry preprint ^1^, we examine the empirical and dynamic behavior of this carrier-set topology. In 1000 Genomes Phase 3, across 156,604,320 SNP pairs from the MHC and NEGR1 regions, pairs on the ∣ *D*^′^ ∣= 1 boundary span a wide range of *r*^2^ and ∣ *C* ∣. Within fixed *r*^2^ strata, *r*^2^ poorly distinguishes nested from non-nested carrier-set configurations, with AUROC values of approximately 0.54–0.62, whereas the boundary-sensitive normalization ∣ *D*^′^ ∣ separates them much more effectively, with AUROC values of approximately 0.90–0.92. The empirical data also obey the predicted *r*^2^ ≤ *p*_*A*_/*p*_*B*_ ceiling.

We then introduce a temporal axis using a two-locus Wright–Fisher model on the same simplex. Carrier-set topology evolves through three motions relative to the ∣ *D*^′^ ∣= 1 boundary: formation or persistence, in which recombination suppression establishes and maintains inclusion without requiring selection; visibility change, in which selection or drift moves *r*^2^ along the boundary while preserving the inclusion relation; and breaking, in which a recombination pulse introduces the previously absent haplotype and dissolves inclusion. A fourth mode, specificity erosion, expands the partner carrier set while preserving inclusion, thereby lowering *P*(*A* ∣ *B*) while keeping *P*(*B* ∣ *A*) and ∣ *D*^′^ ∣ equal to one. This mode shows that asymmetric conditional probabilities are best understood as diagnostic coordinates for carrier-set topology, not as the primary object itself.

Together, these results show that topology and visibility are separable axes of LD structure. Conventional *r*^2^-based scans and carrier-set topology scans therefore answer complementary, not interchangeable, questions.

## 1. Introduction

Linkage disequilibrium (LD) between two biallelic loci is almost always summarized by scalar measures such as *r*^2^and *D*^′^. These measures are useful because they reduce a four-haplotype distribution to a single number describing the strength of allelic association. In practice, *r*^2^is often treated as a measure of tagging quality or detectability: a variant pair with high *r*^2^is visible to an association or imputation procedure, whereas a pair with low *r*^2^is often considered weakly linked or uninformative. This view is appropriate for many tasks, but it compresses a different object that is present in the same haplotype frequencies: the topology of the carrier sets.

For two variants *A*and *B*, the carrier sets *S*_*A*_and *S*_*B*_can be related in several qualitatively distinct ways. One set may be contained within the other, the two may partially overlap, they may be nearly identical, or they may be largely disjoint. These relations are not equivalent to the magnitude of *r*^2^. A rare variant may occur entirely on a common haplotype background, producing exact carrier-set inclusion, while still having low *r*^2^because its allele frequency is much smaller than that of the background variant. Conversely, two balanced variants may have high *r*^2^without representing the same rare–common nesting relation. Thus, LD contains at least two separable axes: the visibility of association to a symmetric scan, and the topology of the carrier sets that generate the association.

This distinction has a natural geometric representation. On the haplotype-frequency simplex, carrier-set inclusion corresponds to a boundary face where one haplotype class is absent. For example, if the haplotype 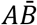 is absent, then every carrier of *A*also carries *B*, so *S*_*A*_ ⊆ *S*_*B*_. The boundary ∣ *D*^′^ ∣= 1is therefore not merely an extreme value of an LD statistic; it is the geometric representation of a set-inclusion relation. In the rare–common regime, this relation is systematically down-weighted by *r*^2^: for a nested rare variant, *r*^2^ ≤ *p*_*A*_/*p*_*B*_, so even complete inclusion cannot reach high *r*^2^when *p*_*A*_ ≪ *p*_*B*_. The information is present in the haplotype distribution, but it is represented poorly by a symmetric Euclidean scalar.

Conditional probabilities provide a convenient way to visualize this topology. *P*(*B* ∣ *A*)measures whether carriers of *A*are also carriers of *B*, whereas *P*(*A* ∣ *B*)measures the converse. These probabilities should not be regarded as the primary biological object; rather, they are directional coordinates for the carrier-set relation itself. Likewise, the decomposition Δ = *P*(*A* ∣ *B*) ™ *P*(*B* ∣ *A*) = *M* + *C*separates marginal-frequency asymmetry from coupling-driven asymmetry ^1,2^, but the central object remains the carrier-set topology on the haplotype simplex.

This reframing is relevant beyond any single locus. Directional LD analyses across the human genome reveal many regions in which strong carrier-set asymmetry coexists with low *r*^2^, including immune clusters, inversions, gene-family arrays, and regulatory regions. The major histocompatibility complex (MHC) is an especially pronounced example: extensive haplotype diversity and long-range LD create many rare–common and partially overlapping carrier-set relations that are poorly summarized by *r*^2^alone. The MHC is therefore not an exception to genome-wide LD structure, but an amplified model system in which the separation between topology and visibility becomes particularly clear.

However, present-day population data provide only a spatial contrast. They show that a marker pair can occupy different carrier-set configurations across ancestries, but they cannot determine how a given configuration was formed, how it persisted, or how it was broken. A positive signed association in one population and a negative one in another ^3,4^may reflect recent selection, admixture, recombination, or simply different present-day projections of an older haplotype architecture. Distinguishing these alternatives requires a controlled temporal axis.

Here we combine empirical analysis of 1000 Genomes Phase 3 data with a two-locus Wright– Fisher model on the same haplotype simplex. We first show that, across 156,604,320 SNP pairs in the MHC and NEGR1 regions, ∣ *D*^′^ ∣= 1boundary pairs span a broad range of *r*^2^, and that *r*^2^cannot distinguish nested from non-nested carrier-set configurations within fixed *r*^2^strata. We then use forward simulation to describe three elementary motions of carrier-set topology relative to the boundary: formation or persistence, visibility change, and breaking. Finally, we distinguish true topology breakdown from specificity erosion, a mode in which one carrier set expands while the inclusion relation itself is preserved. Together, these analyses show that *r*^2^-based LD scans and carrier-set topology scans answer different questions: one asks how visible an association is, whereas the other asks what set relation the association represents.

## 2. Carrier-set topology on the haplotype simplex

For a biallelic pair *A*and *B*, let the four haplotype frequencies be

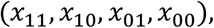

where *x*_11_denotes the frequency of haplotypes carrying both variants, *x*_10_those carrying *A*but not *B, x*_01_those carrying *B*but not *A*, and *x*_00_those carrying neither. The marginal allele frequencies are *p*_*A*_ = *x*_11_ + *x*_10_and *p*_*B*_ = *x*_11_ + *x*_01_.

The carrier sets *S*_*A*_and *S*_*B*_are determined by these same four frequencies. In particular, the absence of one haplotype class corresponds to a qualitative carrier-set relation. If *x*_10_ = 0, then no haplotype carries *A*without *B*, so every carrier of *A*is also a carrier of *B*, and *S*_*A*_ ⊆ *S*_*B*_.

Conversely, if *x*_01_ = 0, then *S*_*B*_ ⊆ *S*_*A*_. Thus, the boundary faces of the haplotype simplex are not only algebraic constraints; they are the geometric representation of carrier-set inclusion.

The conventional statistic *D*^′^detects these boundary faces through normalization by the feasible maximum of *D*, so ∣ *D*^′^ ∣= 1corresponds to the absence of at least one haplotype class. However, ∣ *D*^′^ ∣alone does not specify the full directional interpretation of the relation, and *r*^2^can be small even on the boundary. In the nested case *x*_10_ = 0, the association strength measured by *r*^2^is constrained by

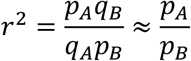

when *p*_*A*_ ≪ *p*_*B*_. Therefore, a rare variant can be perfectly nested within a common background while remaining nearly invisible to *r*^2^.

Conditional probabilities are useful diagnostic coordinates for this topology:

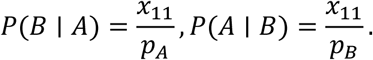

When *P*(*B* ∣ *A*) = 1, carriers of *A*are contained within carriers of *B*. When *P*(*A* ∣ *B*) = 1, the converse holds. When one conditional remains high while the other falls, the inclusion relation and the specificity of the partner set can be separated. These probabilities therefore visualize the carrier-set relation; they are not the primary object of the analysis.

For connection with the Fisher-geometry treatment, the conditional-probability asymmetry can be decomposed as

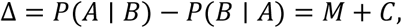

with

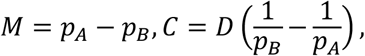

where

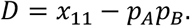

Here *M*is the marginal-frequency contribution and *C*is the coupling-dependent contribution. This decomposition is useful for visualizing directional asymmetry, especially in rare–common pairs, but the central geometric object is the boundary structure of the haplotype simplex and the carrier-set topology it represents.

On the Fisher-information simplex the feasible domain is bounded by the four faces x_h = 0. The face x_10_ = 0 is the nested face: A occurs only on the B background, so the carrier set of A is contained in that of B, |D′| = 1, and P(B|A) = 1 exactly. The three modes developed below are motions of a variant pair toward this face (formation), along it (visibility), and away from it (breaking). The rare–common ceiling follows in two lines: for a nested pair, r^2^ = p_A q_B / (q_A p_B) ≈ p_A / p_B for small frequencies, so a rare variant nested on a common background is bounded away from high r^2^ regardless of phase. The fuller geometric treatment—curvature K(s) = ¼ + (2s™1)^2^/[4s(1™s)], anisotropy, and boundary amplification of det(g)—is given in the geometry preprint (Ichikawa, preprint) ^1^and follows the Wright–Fisher construction of Hofrichter, Jost & Tran (2019) ^5^.

## 3. Empirical structure in 1000 Genomes

### 3.1 Data and methods

Phased haplotypes were taken from the 1000 Genomes Project Phase 3 release ^6^(GRCh37), pooling all 2,504 individuals (5,008 haplotypes) across the five super-populations. Two regions with contrasting LD architecture were examined: the MHC on chromosome 6 (chr6:29.7–33.1 Mb; high haplotype diversity, long-range LD) and the NEGR1 region on chromosome 1 (chr1:71.5–73.0 Mb; typical intergenic structure). SNPs were retained if biallelic with pooled MAF ≥ 0.01. For each retained pair, haplotype frequencies were obtained by direct counting from the phased data (no EM reconstruction), and D, D′, r^2^, |C| = |D|·|1/p_B ™ 1/p_A|, P(B|A), and P(A|B) were computed. Within-region SNP pairs no more than 100 kb apart were enumerated from 58,116 SNPs (52,850 in MHC, 5,266 in NEGR1), giving 156,604,320 pairs in total (154,804,848 in MHC, 1,799,472 in NEGR1). Because C is antisymmetric under exchange of loci, the magnitude |C| is used as the asymmetry measure throughout this section; signed C is used only in the directional analyses of Section 4.

### 3.2 Disagreement among measures at fixed D′ = 1

Among the 156,604,320 pairs, those at D′ = 1 (complete absence of at least one haplotype; n = 21,652,485) span essentially the full r^2^ range (median r^2^ = 0.004; 95.7% of them below r^2^ = 0.2) while |C| runs from 0 to about 0.85 (median 0.078). D′ = 1 is therefore compatible with essentially any combination of r^2^ and |C|, as predicted by their distinct normalizations of D. Two pairs at D′ = 1.00 make the disagreement concrete: a rare–common pair, MAF (0.011, 0.090), with r^2^ = 0.12 and |C| = 0.80; and a balanced pair, MAF (0.264, 0.264), with r^2^ = 1.00 and |C| =0.00. The pairs differ by a factor of roughly nine in r^2^ and, in the limit, unboundedly in |C|—both at the feasible-domain boundary but at very different locations on the curved leaf.

This disagreement is structural rather than a finite-sample artifact: at D′ = 1 the four haplotype frequencies, and hence r^2^ and |C|, are fixed by the two marginal allele frequencies, so the contrast between the two pairs reflects where each sits on the curved leaf and cannot be removed by additional sampling.

### 3.3 Where r^2^ and |C| disagree: a low-r^2^ |C| envelope

Across the full dataset the relationship between r^2^ and |C| is not a simple inverse: their overall rank correlation is in fact weakly positive (Spearman ρ = 0.39 over all pairs, 0.66 among D′ = 1 pairs), because both grow with |D|. What is structural is the envelope. r^2^ = D^2^/(p_A q_A p_B q_B) is maximized when p_A ≈ p_B, whereas |C| = |D|·|1/p_B ™ 1/p_A| is maximized when p_A ≪ p_B or p_A ≫ p_B; the two measures peak in non-overlapping regions of allele-frequency space, so large |C| is reachable only at low r^2^. The upper envelope of |C| descends as r^2^ rises (Fig 1C), and the highest |C| values occur exclusively among low-r^2^, rare-allele pairs, whereas low |C| occurs at every r^2^. The rare–common amplification of |C| is thus a property of this envelope, not a global anti-correlation. Stratifying the D′–r^2^ and D′–|C| relationships by the rarer MAF of each pair shows rare-MAF pairs clustering at low r^2^ and high |C|, and common-MAF pairs at high r^2^ and low |C| (Figure 1; the empirical counterpart of the rare–common regime).

**Figure 1.**
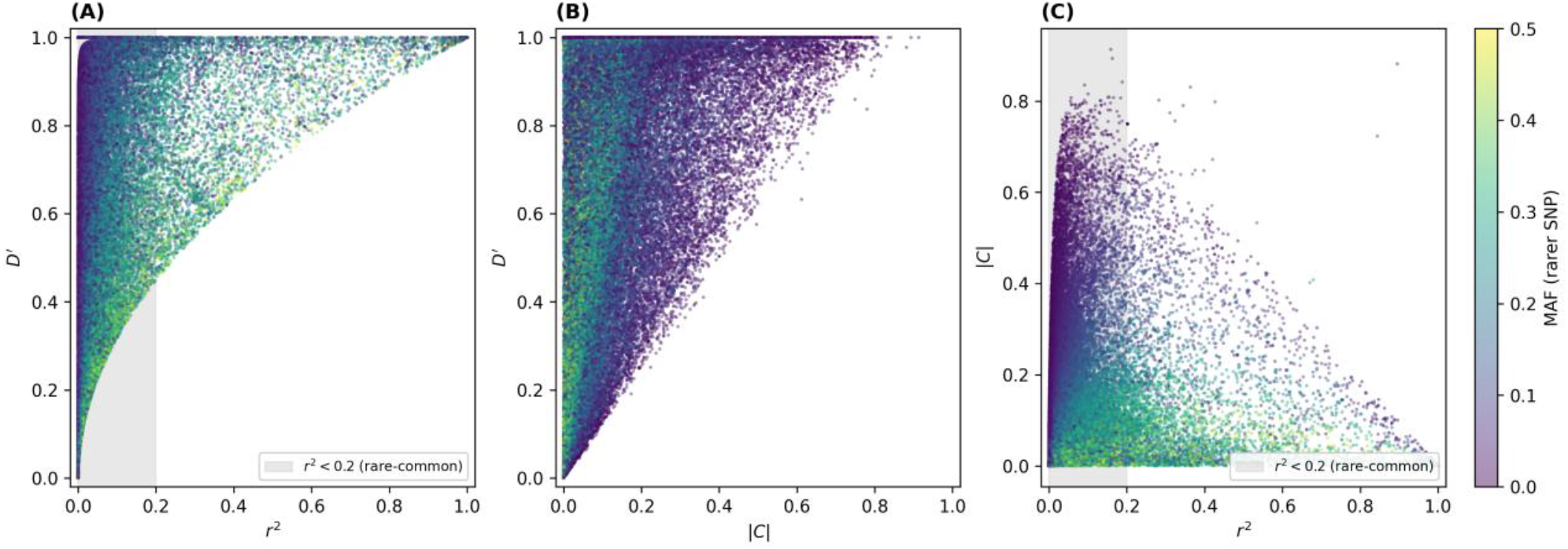
Empirical behavior of LD measures across 156,604,320 SNP pairs (1000 Genomes Phase 3; MHC chr6:29.7–33.1 Mb and NEGR1 chr1:71.5–73.0 Mb; within-region pairs ≤ 100 kb apart, MAF ≥ 0.01; pooled across 2,504 individuals / 5,008 haplotypes). (A) D′ vs r^2^; (B) D′ vs |C|; (C) |C| vs r^2^. Points are colored by the minor allele frequency of the rarer SNP; the shaded band marks the rare–common regime r^2^ < 0.2; a random subsample of 60,000 pairs is shown.

**Figure 2.**
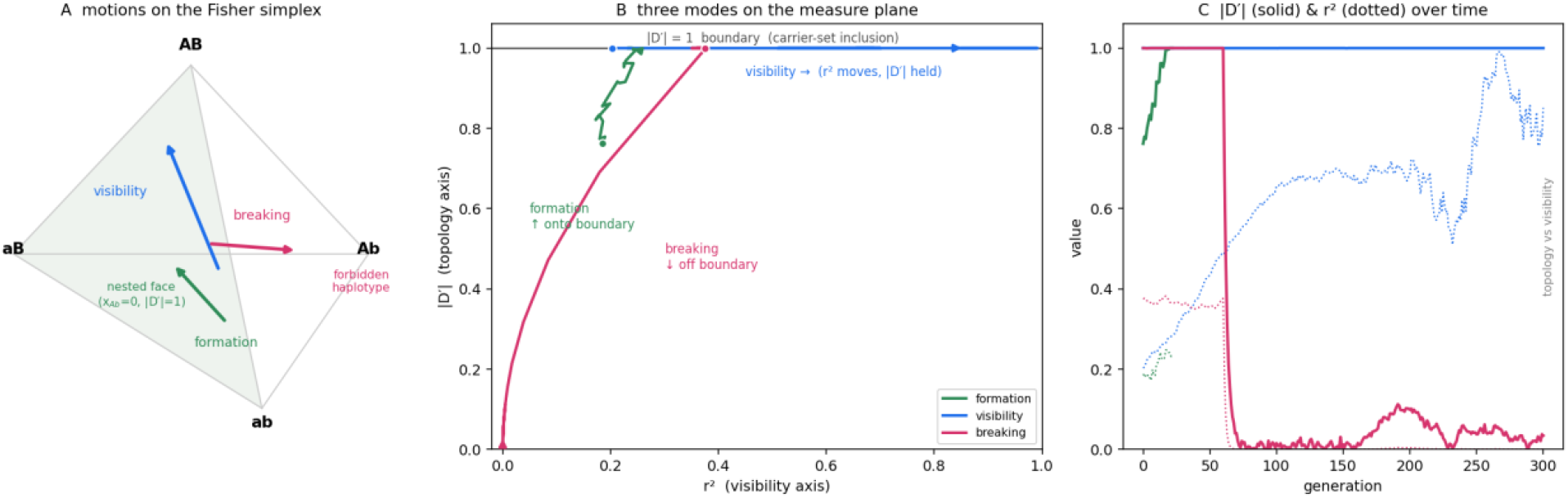
The three evolutionary modes as motions relative to the |D′| = 1 boundary. (A) Fisher haplotype simplex with the nested face shaded and three arrows: formation (interior → face), visibility (along the face), breaking (face → forbidden Ab vertex). (B) The (r^2^, |D′|) measure plane: formation climbs onto the boundary, visibility slides along it (r^2^ moves ≈ 0.2 → 0.85 with |D′| held at 1), breaking drops off it. (C) Time series of |D′| (solid) and r^2^ (dotted) for each mode.

### 3.4 Structural benchmark: r^2^ is blind to nested structure within a fixed r^2^ bin

The sharper prediction is that r^2^, as a Euclidean scalar on a curved manifold, loses information about the partial-order (nested) structure of local haplotypes. For each pair, carrier sets S_A and S_B were computed from phased haplotypes and the pair labeled nested (A → B) if |S_A \ S_B| / |S_A| ≤ 0.05. Pairs were binned by r^2^ (≈ 0.20, 0.50, 0.80), and within each bin the ability of each score to separate nested from non-nested pairs was evaluated by AUROC. Within a fixed bin r^2^ cannot discriminate by construction; the question is whether any other haplotype-frequency scalar carries nested-structure information that r^2^ discards. Results in the MHC region:

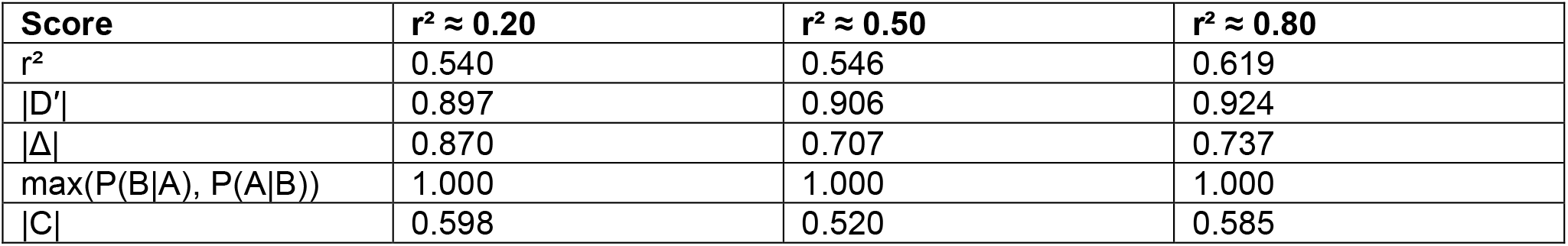

Within each bin r^2^ performs at chance (0.54–0.62), as expected. The conditional-probability score max(P(B|A), P(A|B)) attains AUROC = 1.00, but this is tautological: the nested label |S_A \ S_B|/|S_A| ≤ 0.05 is exactly max(P(B|A), P(A|B)) ≥ 0.95, so this score serves only as a label sanity check, not independent evidence; for the same reason the perfect directional-sign accuracy of sign(Δ) reflects that nesting fixes which allele is rarer (nested ⇒ p_A ≤ p_B ⇒ sign(Δ) = sign(M)) and is not independent evidence. The genuinely non-trivial result is that |D′| retains AUROC ≈ 0.90–0.92 and |Δ| 0.71–0.87: standard frequency-dependent normalizations of D recover nested structure that r^2^ discards. |C| performs weakly here (0.52–0.60), which is expected—|C| isolates the linkage-specific component after removing the frequency-driven part M and is not designed to detect carrier-set inclusion; its value is in directional-sign and rare-variant-amplification tasks (Section 4), not this label.

### 3.5 The structural ceiling r^2^ ≤ p_A / p_B

The ceiling r^2^ ≤ p_A q_B / (q_A p_B) ≈ p_A / p_B predicts that rare–common pairs cannot reach high r^2^ regardless of configuration. This holds in the data: 99.985% of all pairs satisfy the bound, and the few apparent exceedances are numerical (largest excess 3.7×10^−6^), so no pair meaningfully exceeds the ceiling even at D′ = 1; the same rare–common pairs nonetheless reach |C| up to about 0.85 and max(P(B|A), P(A|B)) up to 1.00. The rare–common association signal is present in the data—maximal D′ linkage, exact carrier-set inclusion—but r^2^ geometrically suppresses it while directional normalizations recover it.

## 4. A two-locus Wright–Fisher model: three modes of carrier-set topology

### 4.1 Model

The Fisher manifold of Section 2 is exactly the two-locus haplotype simplex, and its feasible-domain boundary |D′| = 1 is exactly carrier-set inclusion. The minimal setting in which the three modes are well-defined boundary motions is therefore a two-locus, four-haplotype Wright– Fisher process; multi-locus simulation blurs the mapping to the leaf. Each generation updates in the order selection → recombination → mutation → multinomial drift: recombination moves mass by ±rD between coupling and repulsion haplotypes; mutation regenerates absent classes at rate μ (the key knob for formation and breaking); drift is multinomial sampling at 2N = 4,000 (N = 2,000 diploid). The default horizon is T = 720 generations (≈18 kyr at 25 yr/generation). A class at frequency below 1/2N is treated as absent, which assigns each pair a carrier-class (DISJOINT / PARTIAL / STRICT A⊂B / STRICT B⊂A / IDENTICAL). All runs are pre-registered: each possible outcome is assigned a meaning before execution.

### 4.2 Mode 1 — Formation and persistence (recombination-gated)

A new mutation arises on a single background and is therefore nested at birth (|D′| = 1, r^2^ ≈ 0.12); its fate is set by the local recombination rate. At μ = 1e™5, the fraction of surviving pairs that remain strictly nested at gen 720 falls monotonically with r: 0.54, 0.51, 0.46 at r = 0, 1e™5, 1e™4; 0.18 at 1e™3; 0.05 at 1e™2; 0.02 at 0.5, with the crossover near r ≈ 1e™3. Preserved pairs hold |D′| ≈ 1 while r^2^ scatters across a wide range—the rare–common nested configuration of Section 3 generated as a time process—whereas peeled pairs lose both |D′| and r^2^, so the surviving distinction is carried by carrier-class (|D′|), not r^2^. A linkage-equilibrium-start control (interior → boundary genesis) shows the same recombination gating, with face-arrival fraction falling from 0.76 at r = 0 to 0.09 at r = 0.5. Real per-site mutation (≈10^−8^) sits firmly in the recombination-dominated regime; μ = 1e™4 marks the washout boundary.

### 4.3 Mode 2 — Visibility change (selection moves r^2^; topology is invariant)

Starting on the nested face with low recombination, the topology/visibility separation is exact: while A segregates, |D′| and P(B|A) are pinned at 1.0 (on x_10_ = 0, P(B|A) = x_11_/(x_11_+x_10_) = 1 identically), and over the same interval r^2^ swings from 0.12 to 0.75 under selection on the AB thread. Selection thus changes how visible the association is to an r^2^ scan without changing the carrier-set relation. The drift-envelope detection threshold for the selection coefficient is |s| ≈ 0.03 (reps exit the neutral envelope within 13–28 generations for |s| ≥ 0.03, by ≈87 generations for s = +0.01, and essentially never for s = ™0.01), with a characteristic asymmetry favoring positive s. A bottleneck control (N = 100 for 30 generations) inflates the variance of p_A without directional trend and preserves |D′| (0.994 → 0.998).

### 4.4 Mode 3 — Breaking, and the distinction from specificity erosion

A recombination pulse introduces the forbidden haplotype and lifts the pair off the boundary: during the pulse window |D′| drops to 0.45, 0.15, and 0.07 at r_pulse = 1e™2, 1e™1, and 0.5, converting 66–90% of pairs from STRICT to PARTIAL, with a clean dose–duration phase diagram. The break is partially reversible—|D′| recovers from 0.45 toward 0.77 after the pulse— but most pairs retain a PARTIAL trace, so permanent breakdown requires sustained recombination.

A distinct mode must be separated from true breaking. Adding partner-only (aB) haplotypes by admixture lowers P(A|B) while holding |D′| and P(B|A) at 1.0: the carrier set of A stays contained in that of B (STRICT preserved), but the partner set expands, eroding specificity without breaking topology. In the simulation, admixture lowers P(A|B) from 0.73 to 0.10 while the inclusion relation is held (|D′| = 1, P(B|A) = 1): this is specificity erosion, not topology breakdown. The framework resolves the two because they live in different conditionals—a fall in P(B|A) (sensitivity), with the forbidden haplotype appearing and |D′| < 1, is a true break; a fall in P(A|B) (specificity), with |D′| = 1 and P(B|A) = 1 held, is partner-set expansion. This is the decisive case for why both asymmetric conditional probabilities are needed: neither |D′| nor r^2^ can tell the two apart.

### 4.5 The three modes on one canvas

## 5. Discussion

The central object of this study is not conditional-probability asymmetry itself, but the carrier-set topology that such probabilities reveal. Two variants can have carrier sets that are nested, partially overlapping, identical, or largely disjoint, and these relations are encoded in the four-haplotype distribution. Conventional LD measures compress this structure into scalar association strength. As a result, *r*^2^measures how visible an association is to a symmetric LD scan, but it does not determine what kind of carrier-set relation the association represents.

The main result is that topology and visibility are separable axes. In the 1000 Genomes benchmark, many SNP pairs lie on the ∣ *D*^′^ ∣= 1boundary, meaning that at least one haplotype class is absent, yet their *r*^2^values span a broad range. Within fixed *r*^2^strata, *r*^2^provides little information about whether a pair is nested or non-nested, whereas ∣ *D*^′^ ∣and the corresponding conditional coordinates recover the boundary relation. This confirms empirically that the carrier-set topology is present in the haplotype frequencies but compressed out of *r*^2^.

The Wright–Fisher model explains why this separation matters dynamically. Recombination architecture can form or preserve a boundary relation without requiring selection. Selection and drift can then move allele frequencies along the boundary, changing *r*^2^and signed association while preserving the inclusion relation itself. Conversely, recombination events can introduce the previously absent haplotype and lift the pair off the boundary, producing true topology breakdown. In this view, selection usually changes visibility, whereas recombination architecture creates and destroys topology.

The specificity-erosion mode further clarifies why both asymmetric conditional probabilities are needed. If *S*_*A*_remains contained in *S*_*B*_, then *P*(*B* ∣ *A*)and ∣ *D*^′^ ∣can remain equal to one even as *S*_*B*_expands by acquiring many carriers that do not carry *A*. This lowers *P*(*A* ∣ *B*)without breaking the inclusion relation. Thus, a fall in *P*(*B* ∣ *A*)indicates loss of sensitivity and true breaking of the nested relation, whereas a fall in *P*(*A* ∣ *B*)indicates partner-set expansion and specificity erosion. The conditionals are therefore not the headline result; they are the coordinates that distinguish different deformations of carrier-set topology.

This distinction has practical consequences for LD pruning, tagging, rare-variant interpretation, fine-mapping, and imputation reference design. An *r*^2^-based scan asks whether one variant can serve as a statistically visible proxy for another. A carrier-set topology scan asks whether one variant’s carriers are contained within, partially overlap with, or extend beyond another variant’s carriers. These questions are complementary. Treating them as interchangeable risks missing rare–common nested structures that are biologically real but geometrically suppressed by *r*^2^.

The rare–common blind spot is structural, not statistical. The ceiling *r*^2^ ≤ *p*_*A*_/*p*_*B*_follows from the normalization of *r*^2^, not from finite sampling or insufficient power. A rare variant can therefore be perfectly nested within a common carrier background and still remain poorly visible to an *r*^2^-based scan. This is not because the haplotype relation is weak; it is because *r*^2^represents that relation through a frequency-dependent visibility scale. For rare-variant tagging, fine-mapping, and imputation reference selection, reliance on *r*^2^alone therefore imposes a predictable blind spot. Carrier-set-aware measures do not replace *r*^2^, but they recover a different object: the set relation that *r*^2^compresses.

Both directional coordinates are needed because carrier-set topology can deform in more than one way. The specificity-erosion mode in Section 4.4 provides the clearest example. If additional *aB*haplotypes are introduced, the carrier set of *B*expands while the carrier set of *A*remains contained within it. In this case *P*(*B* ∣ *A*)and ∣ *D*^′^ ∣remain equal to one, so the inclusion relation is preserved, but *P*(*A* ∣ *B*)falls because *B*has become a less specific marker for *A*. This is not true topology breakdown. True breaking occurs when the previously absent 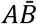 haplotype appears, *P*(*B* ∣ *A*)falls, and the pair leaves the boundary. Thus, *P*(*B* ∣ *A*)diagnoses loss of inclusion, whereas *P*(*A* ∣ *B*)diagnoses expansion of the partner set. The conditionals are not the primary object; they are the directional coordinates needed to distinguish different deformations of the same carrier-set relation.

The unifying object across the related analyses is the ∣ *D*^′^ ∣= 1face of the Fisher haplotype simplex. In the geometric treatment, this face is a curvature-amplified boundary of the feasible domain. In the carrier-set decomposition of tag SNPs, it is the local inclusion relation between variant carrier sets. In the dynamic model developed here, it is the surface toward which variant pairs move during formation, along which they move during visibility change, and away from which they move during breaking. The three papers therefore describe the same geometry from complementary perspectives: why conventional LD measures distort it, where it appears in empirical haplotype data, and how it changes over time.

Several limitations follow directly from this framing. The 1000 Genomes analysis is cross-sectional and therefore spatial: it establishes that carrier-set configurations differ across regions and ancestries, but it cannot date their formation or determine the historical process that produced them. The Wright–Fisher model supplies a controlled temporal axis, but it is deliberately minimal. The clean separation between topology and visibility holds most exactly in the low-recombination regime; with appreciable recombination, strong selection can change visibility and erode topology at the same time. The empirical analysis also emphasizes the MHC, with NEGR1 used as a contrasting non-MHC region, so broader surveys of non-MHC loci are needed to establish how general these modes are across the genome. Finally, the direct empirical temporal test is ancient DNA ^7^: if carrier-set topology is a real dynamic object, its formation, persistence, erosion, and breaking should be observable across time-resolved haplotype data.

The geometry itself is not specific to the MHC or to human immune loci. It is intrinsic to the four-haplotype simplex of any pair of biallelic variants. The human data analyzed here provide an empirical demonstration, and the Wright–Fisher simulations provide a dynamic interpretation, but the underlying distinction is general: *r*^2^measures visibility, whereas carrier-set topology describes the set relation that generates that visibility.

## Methods and implementation

Haplotype frequencies were computed by direct counting from phased 1000 Genomes Phase 3 data; no EM reconstruction was used. Posterior credible intervals (Section 3.2) used a Jeffreys Dirichlet(½, ½, ½, ½) prior on the four haplotype frequencies, updated by observed counts and propagated to r^2^, |C|, and D′ by direct sampling. The forward model is a two-locus, four-haplotype Wright–Fisher process with the per-generation order selection → recombination → mutation → multinomial drift (2N = 4,000; T = 720); observed quantities are D, D′, r^2^, signed-r, P(A|B), P(B|A), Δ = M + C, |C|, M, and carrier-class, with absence thresholded at 1/2N. Random seeds are fixed for reproducibility.

## Data and code availability

1000 Genomes Project Phase 3 data are public. Simulation and analysis code (mode1_formation.py, mode2_visibility.py, mode3_breaking.py, fig2_three_modes.py, and the shared two-locus engine wf_engine.py), together with five supplementary animations (concept, sim_time, sim_mc, region_real, real_mc) and a browser viewer (index.html), are released in the directional-ld-mc repository (https://github.com/mountbook-lab/directional-ld-mc; archived at Zenodo, doi: 10.5281/zenodo.21098987).

